# Automatic antibiotic resistance prediction in *Klebsiella pneumoniae* based on MALDI-TOF mass spectra

**DOI:** 10.1101/2021.10.04.463058

**Authors:** Alejandro Guerrero-López, Carlos Sevilla-Salcedo, Ana Candela, Marta Hernández-García, Emilia Cercenado, Pablo M. Olmos, Rafel Cantón, Patricia Muñoz, Vanessa Gómez-Verdejo, Rosa del Campo, Belén Rodríguez-Sánchez

**Author notes:** Corresponding author, *Email address:* (Alejandro Guerrero-López).

## Abstract

Matrix-Assisted Laser Desorption Ionization Time-Of-Flight (MALDI-TOF) Mass Spectrometry (MS) is a reference method for microbial identification and it can be used to predict Antibiotic Resistance (AR) when combined with artificial intelligence methods. However, current solutions need time-costly preprocessing steps, are difficult to reproduce due to hyperparameter tuning, are hardly interpretable, and do not pay attention to epidemiological differences inherent to data coming from different centres, which can be critical.

We propose using a multi-view heterogeneous Bayesian model (KSSHIBA) for the prediction of AR using MALDI-TOF MS data together with their epidemiological differences. KSSHIBA is the first model that removes the ad-hoc preprocessing steps that work with raw MALDI-TOF data. In addition, due to its Bayesian probabilistic nature, it does not require hyperparameter tuning, provides interpretable results, and allows exploiting local epidemiological differences between data sources. To test the proposal, we used data from 402 *Klebsiella pneumoniae* isolates coming from two different domains and 20 different hospitals located in Spain and Portugal. KSSHIBA outperforms current state-of-the-art approaches in antibiotic susceptibility prediction, obtaining a 0.78 AUC score in Wild Type classification and a 0.90 AUC score in Extended-Spectrum Beta-Lactamases (ESBL)+Carbapenemases (CP)-producers. The proposal consistently removes the need for ad-hoc preprocessing by working with raw MALDI-TOF data, which, in turn, reduces the time needed to obtain the results of the resistance mechanism in microbiological laboratories. The proposed model implementation as well as both data domains are publicly available.

## 1. Introduction

Multidrug-resistant *Klebsiella pneumoniae* is considered a global public health threat by major international health organisations due to its rapid spread, high morbidity, and mortality, as well as the economic burden associated with its treatment and control [1, 2, 3]. Resistance to carbapenems is a major challenge, as recognised by the World Health Organization (WHO) [4], since some carbapenemases can hydrolyse almost all beta-lactam antibiotics. Concretely, over the last two decades, *K. pneumoniae* has shown a great capability to acquire antibiotic-resistant mechanisms, mainly beta-lactamases and carbapenemases. The presence of *K. pneumoniae* isolates hosting these resistance mechanisms complicates the treatment options and the patient’s outcome [5]. Thus, besides the routinely Antimicrobial Susceptibility Testing (AST), rapid diagnostic methods such as MALDI-TOF MS should be implemented in clinical microbiology laboratories for the early detection of multidrug-resistant isolates.

MALDI-TOF MS is designed for microbial identification, but also allows the detection of ESBL and CP due to the different molecular weight of the antibiotic after its hydrolysis by resistant bacteria [6]. This approach is faster than conventional AST (30-60 min vs. 18-24 h), but requires highly trained personnel able to analyse each spectrum profile to detect the AR, so its use is limited to clinical laboratories. However, as suggested in [7], Logistic Regressor (ML) approaches can automatically analyse and predict AR based on the MALDI-TOF MS protein profiles. The most relevant limitation of this methodology is the high complexity of the MALDI-TOF spectrum, which is influenced by the particularities of each lineage and its accessory genome. Hence, two isolates carrying the same CP gene could differ in their MS due to their particular genetic background. Therefore, useful ML models have to be able to jointly learn from their MALDI-TOF MS and their epidemiology to model these particularities.

In the literature, ML has been widely explored for the identification of microbial species. In [8], they identified different clonal lineages of methicillin-resistant *S. aureus* by using ClinProTools [9] black-box private software. This same software was used in [10] to discriminate between contagious and environmental strains of *Streptococcus uberis*. However, other authors prefer to use open-coded models. For example, in [11] the discrimination between *B. anthracis, E. coli, S. pneumoniae 18C-A, and S.pyogenes* based on their MALDI-TOFs was perfectly performed by a Random Forest (RF). Other authors, such as [12], used both an RF and a Support Vector Machine (SVM) to classify different serotypes of Group B streptococcus (GBS). In [13] they also use an RF, a SVM and a multiple Logistic Regressor (LR) to perform strain typing of *S. haemolyticus*.Other approaches, intended for high-dimensional data, such as [14] propose using sparse SVMs to classify the intestinal bacterial composition. However, there is a lack of studies on the prediction of AR using ML, and current works, such as [15], require complex multivariate time series databases to perform AR prediction. Others, such as [16] propose Deep Learning (DL) models which can not work with heterogeneous data. Moreover, current studies that use MALDI-TOF MS require a time-costly preprocessing pipeline commonly performed by using the R package *MALDIquant* (MQ) [17]. After this first step, supervised ML methods such as SVM or RF are applied for the detection of different antibiotic resistances in *S. aureus* [18, 19, 20]. One of the most commonly used tools is the ClinProTools software [9] which contains plug-and-play classification models such as Genetic Algorithm (GA), Supervised Neural Network (SNN), SVM or Quick Classifier (QC) [21], although they are used as a black box. Other previous work dealing specifically with AR prediction in *K. pneumoniae* [22] points out that RF is the best model for predicting the production of CP, however, it was tested over a small dataset (N<100).

Regarding AR prediction, current research is focusing in tree decision predictors, as LightGBM [23] or XGBoost [24]. For example, in [25] they propose to predict AR using a LightGBM, a Multilayer Perceptron (MLP) and an LR. Following same idea, LightGBM has been recently used [26] as a state-of-the-art (SOTA) model to predict AR for *S. aureus*. Another authors [27] also proposed to use LightGBM to predict ceftazidime-resistant *Stenotrophomonas maltophilia*. Then, other researchers [28] propose to use XGBoost as a multi-label predictor for *S. aureus*. However, in [29] they proposed to use classic ML models such as RF, SVM, or LR.

Although supervised SOTA models are a powerful classification tool, they are not designed to deal with high dimensionality data such as the MALDI-TOF MS. Consequently, it is necessary to reduce the dimensionality of these data; in this regard, some authors [30, 31] proposed the use of a GA as a dimensionality reduction technique. Then, they used a SVM as a classifier to perform susceptibility prediction of *S. aureus*. Other approach [32] proposed using a RF as a peak selector to subsequently predict AR using simpler methods such as LR or Linear Discriminant Analysis (LDA). Other authors [33] used unsupervised learning to identify relevant features and then applied Binary Discriminant Analysis (BinDA) [34] and SVM as classifiers. Nowadays, Bayesian models are starting to be applied in this field, as they get rid of cross-validation problems and can provide a predictive distribution with a measure of confidence. A recent study [35] proposed a new SOTA approach where they first reduce the dimensionality of the MALDI-TOF to only 200 peaks per sample by implementing topological peak filtering. Then, they proposed a specific Peak Information KErnel (PIKE) which exploits the correlation between the peaks intensity and their relative position in mass/charge (m/z) dimension. Later, they perform the prediction by a Bayesian probabilistic method, such as a Gaussian Process (GP). Finally, up to our knowledge, any SOTA work tackles the epidemiological differences of the MALDI-TOF spectra. Following this new approach, the same authors [36] proposed to use hierarchical clustering to induce phylogenetic structure and then use again their GP-PIKE model.

As seen in the literature, all SOTA models [9, 30, 19, 18, 31, 20, 33, 22, 37] tend to share a common pipeline: (1) a time-costly MALDI-TOF MS data preprocessing, (2) a dimensionality reduction technique, and (3) a cross-validation step to choose the model hyperparameters. In this paper, we propose to simplify this pipeline while providing explainable results, including epidemiological information and outperforming SOTA models in AR prediction. Then, using KSSHIBA as a predictor for AR is motivated by three main factors: it can work directly with raw data, meaning that do not need preprocessing steps; KSSHIBA performs dimensionality reduction by kernel methods, as some current SOTA models do, such as SVMs or GPs; lastly, due to its Bayesian nature, it automatically deals with parameter selection by optimisation. Furthermore, KSSHIBA brings interpretable solutions as it explains each of the views by a linear product with a matrix weight that changes across views, enabling us to identify which latent dimensions are explaining each of the views. For this purpose, we adapt KSSHIBA to efficiently deal with MALDI-TOF MS data by combining it with kernel functions equivalent to PIKE [35]. This way, we get to simplify the standard pipeline by:

- **Avoiding external preprocessing** since KSSHIBA can efficiently handle raw MALDI-TOF MS data, getting rid of time-costly MQ preprocessing.
- **Performing a double dimensionality reduction**, firstly, through the kernelised data representations, which uses matrices of dimension number of samples (*N*) instead of the number of features (*D*). And, secondly, obtaining a common low-dimensional latent representation of all input data sources.
- **Eliminating hyperparameter tuning**, since the Bayesian nature of the model, entails the ability to automatically optimise the model parameters by maximising the variational lower bound.

Moreover, we **tackle the epidemiological differences** by using its multiview nature, indicating the origin centre of each spectrum. All this while providing **interpretable results** since it calculates a weight matrix for each view, capable of explaining how they correlate.

To endorse the proposal, we use two different epidemiological domains of *K. pneumoniae* isolates: one collection of 282 samples coming from Hospital General Universitario Gregorio Marañón (GM), and another collection of 120 samples coming from Hospital Universitario Ramón y Cajal (RyC). Over these two data collections KSSHIBA outperforms current SOTA models, such as a GP, an SVM, a LightGBM, an MLP, and an XGBoost. Namely, KSSHIBA scores 0.78 and 0.90 AUC scores in Wild Type (WT) and ESBL+CP, respectively, in the GM collection. Then, KSSHIBA scores 0.70 and 0.68 AUC score in WT and ESBL+CP, respectively in the RyC collection.

The article is organised as follows: Section 2 presents the *K. pneumoniae* collections, the technical adaptations to kernelised Sparse Semi-Supervised Interbattery Bayesian Analysis (KSSHIBA), and the models that were chosen to compare KSSHIBA to. Section 3 presents two experiment setups, intra-domain and inter-domain, where the prediction of ESBL and CP is performed, and the latent space explainability. Finally, in Section 4 a final conclusion is stated, and the main results are highlighted.

## 2. Materials and Methods

### 2.1. Isolates selection and processing

We include two different data domains. The first data domain has 282 consecutive clinical *K. pneumoniae* isolates collected between 2014 and 2019 and later isolated at Hospital General Universitario Gregorio Marañón (GM). Therefore, this data domain is called from now on the GM domain. The second data domain has 120 isolates which were characterised in surveillance programs (STEP and SUPERIOR) [38, 39] sourcing from 8 Spanish and 11 Portuguese hospitals. The AST determination of these 120 isolates has been performed in the Hospital Universitario Ramón y Cajal (RyC). Therefore, this data domain is called from now on the RyC domain.

The AST determination has been performed separately in their origin centre by the automated broth microdilution method Microscan® System (Beckman-Coulter, CA, USA) using EUCAST (2021) common criteria. The presence of ESBL/CP genetic-resistant mechanisms has been corroborated by molecular testing. Each isolated is labeled as WT, ESBL-producers or ESBL+CP-producers as shown in Table 1.

**Table 1:** Dataset detailed by domain and label types.

| Dataset | Label | Samples |
| --- | --- | --- |
| GM | WT | 85 |
|  | ESBL | 6 |
|  | ESBL+CP | 191 |
| RyC | WT | 9 |
|  | ESBL | 58 |
|  | ESBL+CP | 53 |

Isolates have been kept frozen at −80°C in skimmed milk and, after thawing, they have been cultured overnight at 37°C in Columbia Blood agar (bioMérieux, Lyon, France) during 3 subcultures for metabolic activation. The MS analysis has been centralised and performed by the same operator using an MBT Smart MALDI Biotyper mass spectrometer (Bruker Daltonics, Bremen), in 6 separated replicates (2 positions on 3 consecutive days). The protein extraction has been performed by adding 1μl 100% formic acid and then drying at room temperature. Next, 1μl of HCCA matrix solution (Bruker Daltonics) has been added to each spot. The MALDI-TOF spectra has been acquired in positive linear mode in the range of 2 kDa to 20 kDa, using default settings [40], although only data between 2,000-12,000 m/z [41, 42] has been used.

The Ethics Committees of both GM and RyC (codes MICRO.HGUGM.2020-002, and 087-16, respectively) have approved this study. The study has been performed on microbiological samples, not human products, and patient-informed consent has not been required.

### 2.2. Multiview KSSHIBA for MALDI-TOF MS data

In this context, we propose to adapt KSSHIBA [43] model, the kernelised version of SSHIBA [44], to deal with MALDI-TOF MS data so that we can efficiently predict the CP and ESBL susceptibility for each isolate. KSSHIBA is a Bayesian multiview semi-supervised model intended to deal with high-dimensional data because, on the one hand, works in the data space (*N* × *N*) being *N* << *D* by means of using kernel data representations, and, on the other hand, it projects all input views to a common low-dimensional latent space.

To tailor the model to the data particularities, we propose to use KSSHIBA to solve a multi-view problem composed of 3 views: one view is the MALDI-TOF MS data kernelised, another view is the AR we want to predict, and another is the domain label representing the hospital where the data belong. The graphical model is shown in Fig. 1^1^. By this multiview scheme, we are able to tackle the different particularities of MALDI-TOF MS data. First, the kernelised view efficiently deals with the high-dimensionality particularity of MALDI-TOF MS data. Secondly, by adding the domain view, we particularly handle possible epidemiological differences between isolates. Finally, by having a low-dimensional common space, we model the interaction between the domain, the data, and its AR providing explainable results.

**Figure 1:**
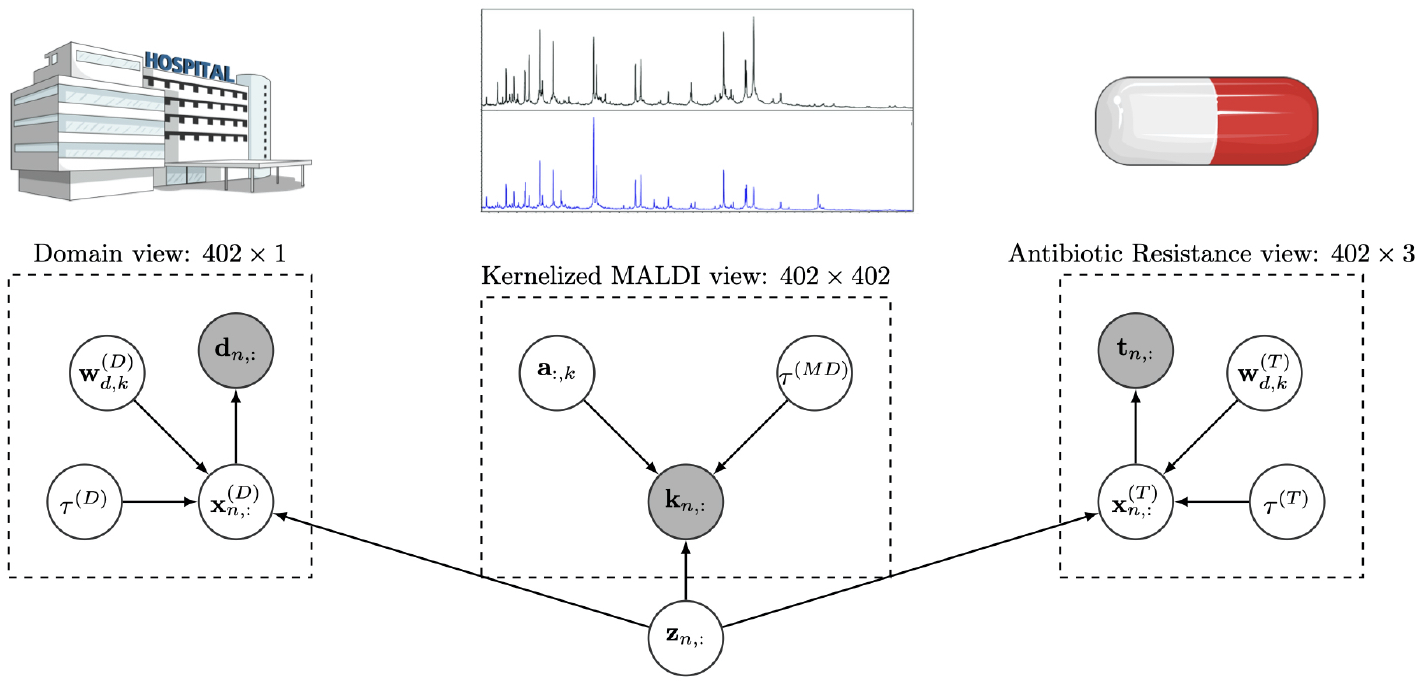
Probabilistic graphical model for the evaluated data set: view **D** corresponds to the label of the domain they come from (GM or RyC), view **M** corresponds to the kernelised MALDI-TOF MS data, and view **T** corresponds to the AR (WT, ESBL or ESBL+CP). The white circles represent random variables that the model learns, while the grey circles represent the observations.

The MALDI-TOF data, i.e. the first view, is kernelised to reduce its dimension to 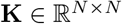, where *N* is the number of samples in each experiment and *N* << *D*. Moreover, choosing different kernel functions allows modelling the relations between peaks in different ways, such as using PIKE or Radial Basis Function (RBF) kernels. Then, each row of **K** represents a kernelised observation, denoted as **k**_n,:_:

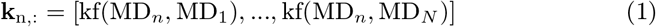

where kf(MD_*a*_, MD_*b*_) is a kernel function between MD_*a*_ and MD_*b*_, which are an arbitrary pair of MALDI-TOF mass spectra.

In this view, KSSHIBA considers that a common low-dimensional latent variable vector 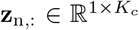 is linearly combined with a set of dual variables 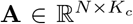, where *K_c_* is the dimension of the low-dimensional latent space, and a zero-mean Gaussian noise *τ*^(M)^ to generate each row of the kernelised observations **k**_n,:_, as:

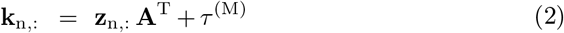

where the prior over the latent space is given by 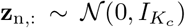 and the prior over the *k*-th column of the dual variables **A**, 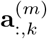, is given by 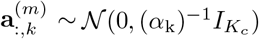. Thus, the random variable α_k_ ~ Γ(*a, b*) follows an Automatic Relevance Determination (ARD) prior [45] over the columns of **A** to automatically select the columns of **z**_n,:_ that are indeed relevant to explain the current data view.

For the AR observations, denoted as T, we propose to use a one-hot encoding for the WT, ESBL and ESBL + Carbapenemases (ESBL+CP) tags. Likewise, for the data domain, denoted as *D*, we consider binary encoding where a 0 value means that the data come from the GM domain and a value of 1 means that the data come from the RyC domain.

To accommodate for these two binary observations, we first consider that there exist two real latent variables **X**^(m)^, *m* ∈ {T, *D*}, that are generated by the common low-dimensional latent variable 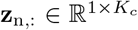, and, then, are linearly combined with a projection matrix 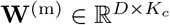 (where *D* is the observation dimension) and a zero-mean Gaussian noise *τ*^(m)^, as follows:

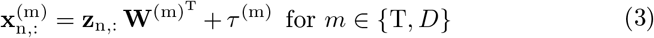

where **W**^(m)^’s prior is identical to **A**’s to automatically select which columns of **z**_n,:_ are needed to explain these two views. Then, we are able to generate **T**^(m)^ by conditioning to this new latent representation **X**^(m)^ using an independent Bernoulli probability model [46], for AR view, as:

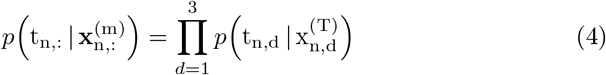

where:

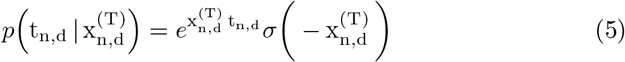

and for domain view, as it is binary:

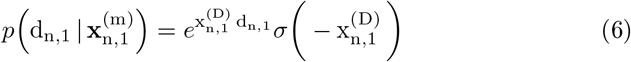

The model is trained by evaluating the posterior distribution of all the random variables posteriors given the observed data. These posteriors are approximated through mean-field variational inference [47] maximising the Evidence Lower BOund (ELBO). For more details, see [43, 44]. Furthermore, the Bayesian nature of the model allows it to work in a semi-supervised fashion, using all available information to determine the approximate distribution of the variables. In turn, the model can marginalise out any type of missing values in the data, as well as predict the test samples for AR by sampling its variational distribution. Table 2 shows the mean-field factor updated rules for each random variable in the KSSHIBA model.

**Table 2:**
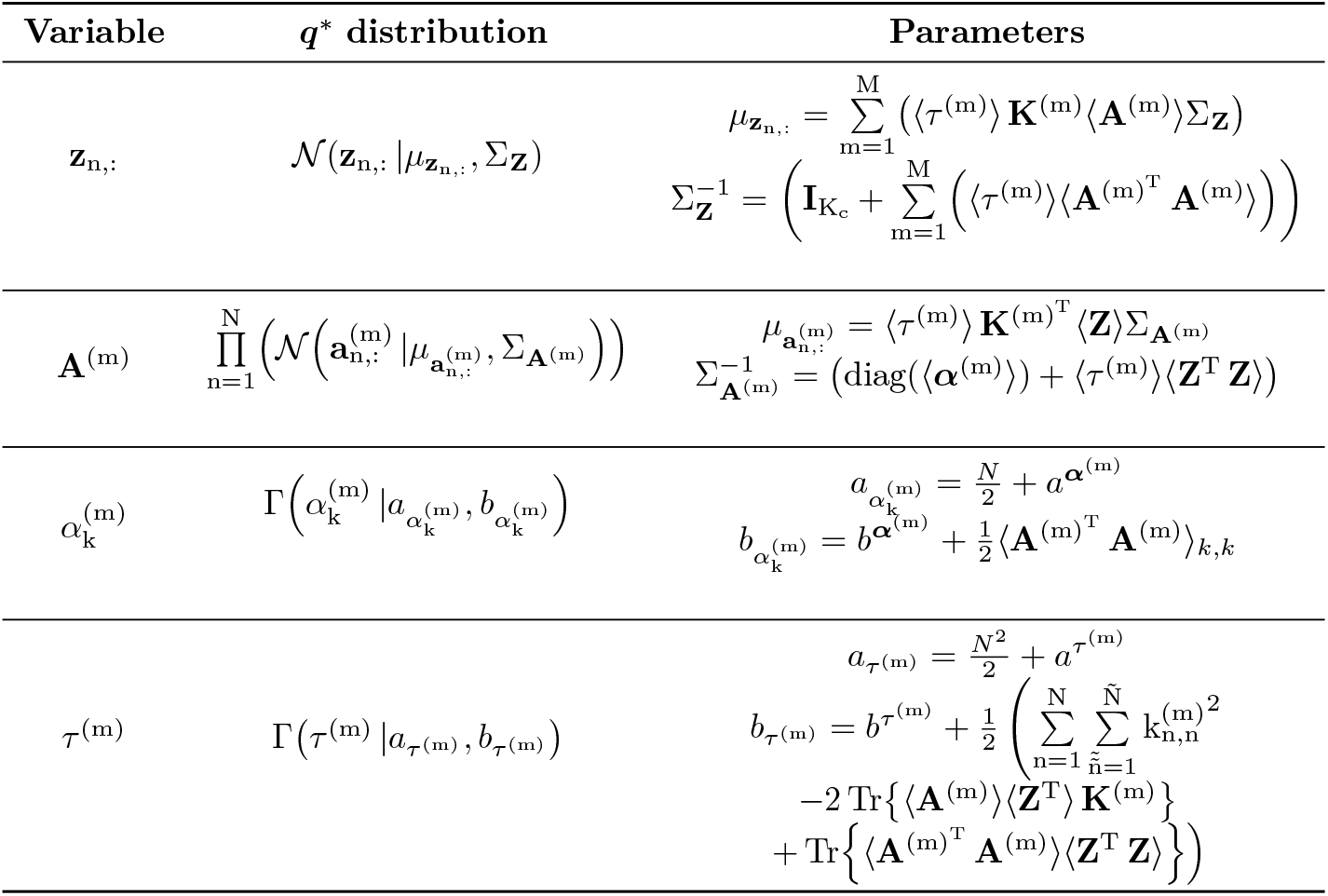
Updated rules, obtained by a mean field approximation, of *q* distribution for the different variables of the KSSHIBA model.

In short, KSSHIBA is a hierarchical model which projects all heterogeneous input views into a common latent space in a linear way. Then, each prediction can be understood as a Bayesian Neural Network (BNN) whose input is determined by the common latent space **z**_n,:_. Finally, the complexity of training this model is the inverse computation of 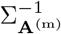 and 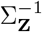 which are matrices of size *K_c_ × K_c_* being *K_c_* the latent space dimension, which is automatically pruned. In particular, we used *K_c_* < 100 for all the experiments in this article.

The implementation of KSSHIBA and all experiments found in this article are publicly available^2^.

### 2.3. Model training and validation

We study two different scenarios: (1) training and testing in each domain separately (intra-domain analysis), and (2) training and testing in both domains simultaneously (inter-domain analysis). For each analysis, we divide each domain into 5 random training-test folds. Due to the label imbalance seen in Table 1, we correct it in each training fold by oversampling the minority class, which ultimately results in stratified folds with a consistent class ratio.

For inter-domain analysis, we merge the two randomised 5-folds of training previously used in the first analysis in two different ways: (1) directly combining both domains, i.e., training with only two views (kernelised MALDI-TOF view and AR view) and (2) combining both domains by adding a third view indicating to which domain each data belongs. In other words, in the first case, we do not use the *D* observations, meanwhile, in the second case, we use the *D* observations. In this way, we analyse the importance of knowing the data origin.

Finally, the performance is measured in terms of Area Under the ROC Curve (AUC) of AR prediction over the test folds.

### 2.4. Methods under study

We compare KSSHIBA with a SVM and a GP since all three models can work with kernel formulations. As a kernel function, we first test a nonlinear approach, such as RBF [48] that is given by:

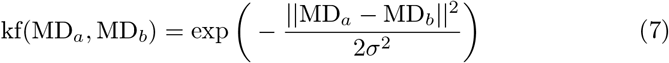

where *σ* is the variance hyperparameter. Then, we also test a linear kernel [49] which follows:

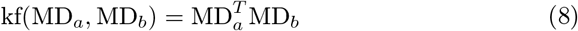

In both cases, MD_*a*_ and MD_*b*_ are a pair of MALDI-TOF spectra.

Finally, we work with a SOTA kernel function called PIKE [35], which exploits the nonlinear correlations between the MALDI-TOF peaks as follows:

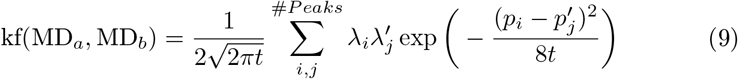

where *t* is a smoothing parameter that has to be cross-validated, λ_*i*_, λ_j_ correspond to the intensity value of each pair of peaks, and *p_i_, p_j_* is their m/z position in the spectra. Let’s remind that each MALDI-TOF consists of 12,000 different peaks. Due to the computational cost to evaluate Eq. 9 in that amount of peaks, the spectra are preprocessed beforehand by topological peak selection [35]. This peak selection is a simple peak detection method based on the persistence concept from computational topology which automatically results in a peak detection because local maxima exhibit high persistence values by construction in MALDI-TOF data. Following the indications of the authors of [35], only 200 peaks per sample are kept.

Since we are solving a multi-class classification problem, we train the SVMs and GPs in a one-vs-all scheme. Besides, we also compare ourselves against a multitask RF.

Regarding cross-validation, we use an inner 5-fold over the training folds to validate all hyperparameters. To do so, we followed a 5-fold grid search cross-validation technique to cross-validate the parameter C in the set of values {0.01, 0.1, 1, 10} for the SVM, and for the RF we adjust the number of estimators and the maximum number of features exploring the values {50, 100, 150} and {*auto, log*2}, respectively. For both KSSHIBA and GP, the hyperparameters are optimised by maximising the ELBO and the marginal log-likelihood of the data, respectively. When using PIKE, we also cross-validate t smoothing value in the range of {1, 5, 10}.

Finally, to demonstrate that KSSHIBA does not need any external preprocessing, we use our model with and without MQ preprocessing. When we use MQ we denote it using the prefix *MQ*-, e.g., MQ-KSSHIBA.

## 3. Results

In this section, we present the results obtained using the proposed model and the different SOTA algorithms. First, we study the classification performance in the intra-domain scenario. Later, we analyse the performance in the inter-domain scenario to evaluate the advantages of working with multi-view data sources. Finally, we study the latent space projection learned by KSSHIBA to understand the correlation between the source domain and the labels.

### 3.1. Intra-domain scenario

Table 3 and Table 4 summarise the results obtained by training and testing independent models for each domain (GM and RyC). The name of every model is constructed by three terms, indicating *Preprocessing-Model-Kernel*. For example, KSSHIBA-RBF means that we use raw MALDI-TOF data and the KSSHIBA model with an RBF kernel function. In contrast, MQ-GP-PIKE means that we use the MQ package to preprocess MALDI-TOF data and a GP model with the PIKE kernel.

**Table 3:** Results of non-linear models in the intra-domain scenario in terms of mean AUC and standard deviation w.r.t. the 5 random splits. The best result for each case is shown in bold. The last row indicates the weighted Average Performance over all the data.

| Dataset | Label | KSSHIBA<br>RBF | KSSHIBA<br>PIKE | MQ-GP<br>PIKE [35] | SVM<br>RBF | RF |
| --- | --- | --- | --- | --- | --- | --- |
| GM | WT | 0.61±0.14 | 0.71±0.16 | **0.75±0.11** | 0.67±0.12 | 0.70±0.17 |
|  | ESBL | **0.57±0.28** | 0.56±0.32 | 0.35±0.14 | 0.40±0.29 | 0.39±0.21 |
|  | ESBL+CP | **0.85±0.14** | 0.78±0.09 | 0.79±0.07 | 0.82±0.19 | 0.80±0.19 |
| RyC | WT | 0.47±0.35 | **0.64±0.19** | 0.56±0.20 | 0.45±0.15 | 0.57±0.26 |
|  | ESBL | 0.70±0.10 | 0.43±0.09 | 0.43±0.11 | **0.72±0.14** | 0.69±0.10 |
|  | ESBL+CP | 0.67±0.12 | 0.43±0.09 | 0.55±0.05 | 0.71±0.17 | **0.71±0.07** |
| Average Performance |  | **0.74 ± 0.13** | 0.66 ± 0.18 | 0.68 ± 0.17 | **0.74 ± 0.11** | 0.71 ± 0.11 |

**Table 4:** Results of linear models in the intra-domain scenario in terms of mean AUC and standard deviation w.r.t. the 5 random splits. The last row indicates the weighted Average Performance over all the data.

| Dataset | Label | KSSHIBA<br>LINEAR | GP<br>LINEAR |
| --- | --- | --- | --- |
| GM | WT | 0.70±0.15 | 0.70±0.18 |
|  | ESBL | 0.46±0.19 | 0.54±0.18 |
|  | ESBL+CP | 0.77±0.16 | 0.80±0.20 |
| RyC | WT | 0.49±0.22 | 0.48±0.28 |
|  | ESBL | 0.59±0.08 | 0.58±0.14 |
|  | ESBL+CP | 0.66±0.05 | 0.62±0.06 |
| Average Performance |  | 0.70 ± 0.09 | **0.71 ± 0.11** |

For the GM domain, as shown in Table 3, KSSHIBA outperforms the baselines in terms of AUC in both ESBL and ESBL+CP prediction. Nonlinear kernels provide the best results in the three tasks, as shown in the comparison between Table 3 and Table 4. Specifically, the RBF kernel is the best choice for both ESBL and ESBL+CP while the PIKE kernel outperforms all other functions in WT prediction.

In contrast, the RyC domain turns out to be the most complex to deal with, since none model works correctly over the three labels simultaneously. Despite this, we again observe that nonlinear techniques such as PIKE, RBF, or RF performed better than linear ones.

### 3.2. Inter-domain scenario

Table 5 and Table 6 show the results obtained by linear and non-linear models, respectively, when trained jointly on both GM and RyC domains. For this scenario, in the case of using the multiview nature of KSSHIBA, it is indicated by *Preprocessing-Model Kernel Domain*. For example, KSSHIBA-LINEAR-DOMAIN means that there is no preprocessing, the KSSHIBA model is used with linear kernel function, and we have added the domain labels.

**Table 5:** Results of linear models in the inter-domain scenario in terms of mean AUC and standard deviation w.r.t the 5 random splits. The best result for every case is shown in bold. The last row indicates the weighted Average Performance over all the data.

| Dataset | Label | KSSHIBA<br>LINEAR<br>DOMAIN | MQ-KSSHIBA<br>LINEAR<br>DOMAIN | KSSHIBA<br>LINEAR | GP<br>LINEAR | SVM<br>LINEAR |
| --- | --- | --- | --- | --- | --- | --- |
| GM | WT | **0.77±0.11** | **0.78±0.06** | 0.72±0.14 | 0.76±0.10 | 0.62±0.13 |
|  | ESBL | 0.46±0.19 | 0.51±0.25 | 0.39±0.21 | 0.43±0.20 | 0.39±0.21 |
|  | ESBL+CP | **0.88±0.08** | **0.90±0.05** | 0.86±0.08 | 0.86±0.08 | 0.85±0.08 |
| RyC | WT | **0.70±0.16** | 0.48±0.28 | 0.66±0.16 | 0.68±0.17 | 0.59±0.20 |
|  | ESBL | 0.55±0.09 | 0.53±0.07 | 0.49±0.09 | 0.60±0.10 | **0.69±0.12** |
|  | ESBL+CP | **0.68±0.10** | 0.63±0.10 | 0.64±0.06 | 0.64±0.04 | 0.66±0.14 |
| Average performance |  | **0.77 ± 0.13** | **0.77 ± 0.18** | 0.74 ± 0.17 | 0.76 ± 0.13 | 0.74 ± 0.13 |

**Table 6:** Results of non-linear models in the inter-domain scenario in terms of mean AUC and standard deviation w.r.t the 5 random splits. The MG-GP PIKE means reproducing the work done in [35]. The last row indicates the weighted Average Performance over all the data.

| Dataset | Label | KSSHIBA<br>PIKE<br>DOMAIN | KSSHIBA<br>RBF<br>DOMAIN | MQ-GP<br>PIKE [35] | SVM<br>RBF |
| --- | --- | --- | --- | --- | --- |
| GM | WT | 0.64±0.17 | 0.59±0.16 | **0.74±0.14** | 0.65±0.13 |
|  | ESBL | **0.44±0.37** | 0.32±0.23 | 0.40±0.12 | 0.40±0.19 |
|  | ESBL+CP | 0.73±0.12 | 0.81±0.12 | 0.77±0.10 | **0.85±0.08** |
| RyC | WT | 0.51±0.14 | **0.63±0.07** | 0.62±0.22 | 0.57±0.26 |
|  | ESBL | 0.39±0.14 | 0.66±0.05 | 0.49±0.09 | **0.69±0.12** |
|  | ESBL+CP | 0.51±0.14 | 0.59±0.11 | 0.59±0.06 | **0.68±0.14** |
| Average performance |  | 0.62 ± 0.15 | 0.70 ± 0.13 | 0.69 ± 0.13 | **0.75 ± 0.12** |

When combining both domains, the linear version of KSSHIBA DOMAIN outperforms all SOTA models for both WT and ESBL+CP predictions, as shown in Table 5. Besides, KSSHIBA DOMAIN LINEAR also outperforms all models in the previous experiment which were targeted to each particular domain. Since KSSHIBA works with heterogeneous data, such as the domain label, it can exploit the information inherent in both data sets.

Regarding the kernel function, when using the RBF kernel, as seen in Table 6, the results worsen significantly making it clear that both distributions are far apart and, therefore, a linear kernel can better explain these differences. This is because setting a common *γ* parameter in the RBF kernel that adequately explains the similarity of the data in both domains is not feasible. Thus, a simpler linear kernel gets better results, as shown in Table 5.

If we compare the versions of KSSHIBA with domain view and without it, e.g. KSSHIBA LINEAR DOMAIN versus KSSHIBA LINEAR in Table. 5, the improvement in the first case indicates that the model can get rid of the possible local epidemiological bias induced by the domain, and properly merges both datasets.

In addition, KSSHIBA shows a new advantage in AR prediction: no external preprocessing with MQ is required. Although MQ preprocessing presents better results in the GM domain, it performs poorly in the RyC domain. The MQ preprocessing pipeline defines reference peaks based on the most frequent peaks. We hypothesise that this pipeline leads the data to be biased to the bigger domain, being GM in our scenario. As we can see, using the raw data allows us to maintain an almost identical performance in the GM domain while we improve all predictions for the RyC domain.

Regarding the rest of the models, the comparison between Table 5 and Table 6 leads to the fact that linear kernels are better for the prediction of both WT and ESBL+CP. This indicates that in the first experiment, by only taking into account one domain, the model was overfitted. Whereas, by adding out-of-distribution data, the linear kernel can generalise better.

As seen in the literature review, current research over AR prediction is done by using LightGBM, XGBoost and MLP neural networks. Therefore, in Table 7 we compare them. Both XGBoost and MLP perform promisingly good. However, none of them outperforms KSSHIBA in Average performance score nor provides decent results in the RyC dataset tending to overfit, as the other models, to the overpopulated dataset.

**Table 7:**
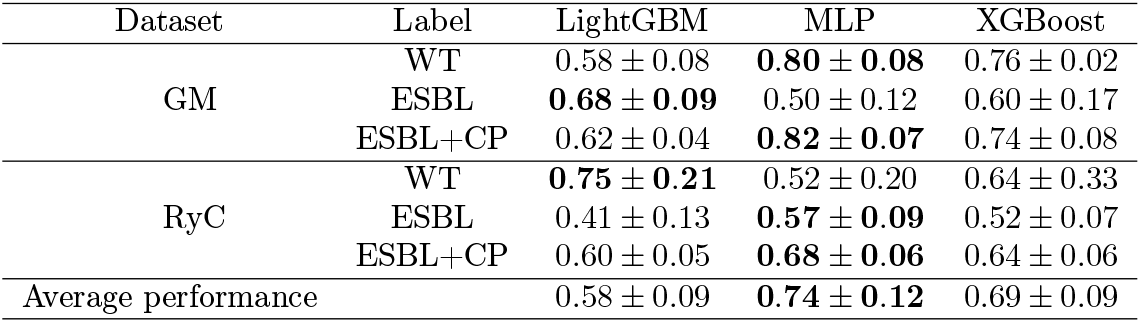
Results of SOTA not based on kernel methods in terms of mean AUC and standard deviation w.r.t the 5 random splits. The last row indicates the weighted Average Performance over all the data.

| Dataset | Label | LightGBM | MLP | XGBoost |
| --- | --- | --- | --- | --- |
| GM | WT | $0.58 \pm 0.08$ | **$0.80 \pm 0.08$** | $0.76 \pm 0.02$ |
| | ESBL | **$0.68 \pm 0.09$** | $0.50 \pm 0.12$ | $0.60 \pm 0.17$ |
| | ESBL+CP | $0.62 \pm 0.04$ | **$0.82 \pm 0.07$** | $0.74 \pm 0.08$ |
| RyC | WT | **$0.75 \pm 0.21$** | $0.52 \pm 0.20$ | $0.64 \pm 0.33$ |
| | ESBL | $0.41 \pm 0.13$ | **$0.57 \pm 0.09$** | $0.52 \pm 0.07$ |
| | ESBL+CP | $0.60 \pm 0.05$ | **$0.68 \pm 0.06$** | $0.64 \pm 0.06$ |
| Average performance | | $0.58 \pm 0.09$ | **$0.74 \pm 0.12$** | $0.69 \pm 0.09$ |

Regarding ESBL prediction, no model achieves high AUC values over one task without losing the remaining predictions. We hypothesise that there is not enough data about ESBL to properly learn all the variability shown in this AR profile. ESBL, as shown in Table 1, only presents 64 samples being the AR profile less represented. Therefore, we would need more data from this AR to be able to generalise correctly.

### 3.3. Latent space analysis

Since KSSHIBA LINEAR DOMAIN has exhibited the best performance, we here analyse the latent space projection learned to understand the importance of the domain label.

Fig. 2 represents the average weight of each latent factor **W**^(m)^ for *m* ∈ {MD, T, *D*}, i.e. the average across every column. We recover **W**^(MD)^ by moving from the dual space **A** variables to primal space as follows:

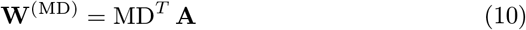

where MD are the raw MALDI-TOF observations. Due to the sparsity induced by the ARD prior explained above, **W**^(m)^ automatically selects the *k* features of **z**_n,:_ that are only relevant for each m data view.

**Figure 2:**
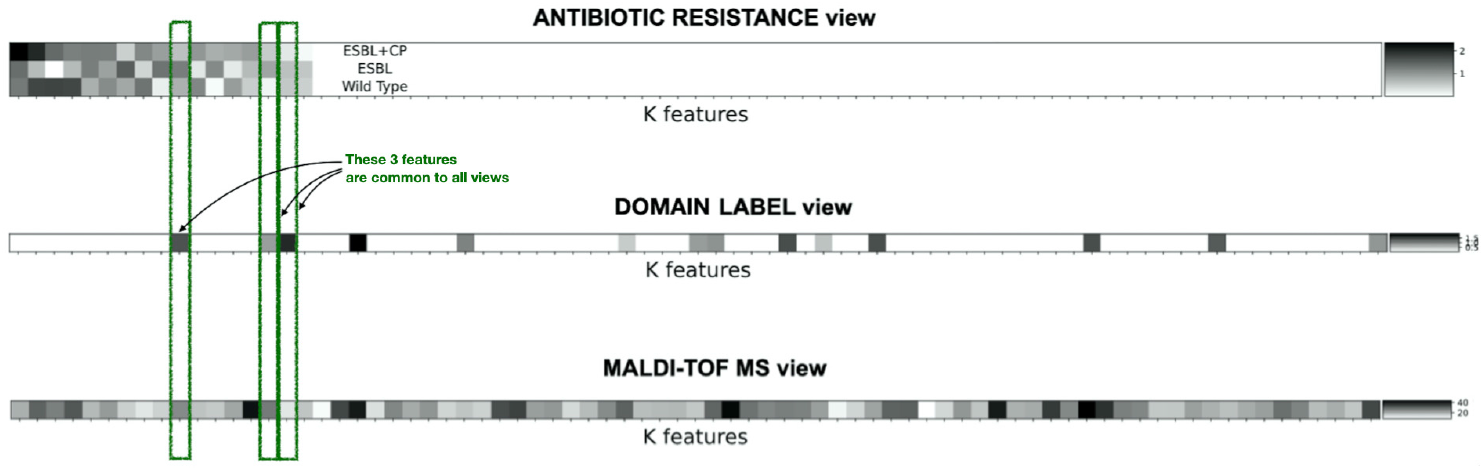
Latent space correlation between input views. Each row represents the mean of each 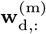 *d*-row having then 76 values, one per each *k* latent feature. Each subplot represents one **W**^(m)^ matrix per view. The most important features (the highest weight value) are represented in black, and the least important features (the lowest weight value) are represented in white. Finally, the features were ordered by their relevance to the prediction task

In this case, KSSHIBA decided that only 76 latent features are needed, as shown in Fig. 2. It is noteworthy that, from these 76 variables, only 14 latent factors are used to predict the label AR. From these 14, all of them share information with the MALDI-TOF view, but only 3 of them simultaneously correlate all available information. Finally, note that 51 private latent features are necessary for the MALDI-TOF view, which corresponds to an unsupervised projection of the data that exclusively models the behaviour of the MALDI-TOF view, as a Principal Component Analysis could do.

Fig. 2 also shows that there is a correlation between the original domain of each strain and its AR, as they have 3 shared latent features. In addition, the domain label is also used to explain the projection of the MS, proving that the MALDI-TOF distributions differ based on epidemiology.

Regarding the low dimension projection, in Fig. 3 we use a T-distributed Stochastic Neighbor Embedding (t-SNE) to project to a 2-dimensional space the 14 latent features that are relevant to explain the domain view. Both domains follow two different distributions, and a simple linear classifier can separate them. The GM domain presents a more compact space as they are all coming from the same hospital. In contrast, as the RyC domain is a collection of 19 different hospitals, we can see that it presents a sparser distribution with different clusters of data points. The four black circles in Fig. 3 correspond to four different hospitals of the RyC domain collection. This means that our model clusters the data not only by domain but also by the hospital without knowing that information.

**Figure 3:**
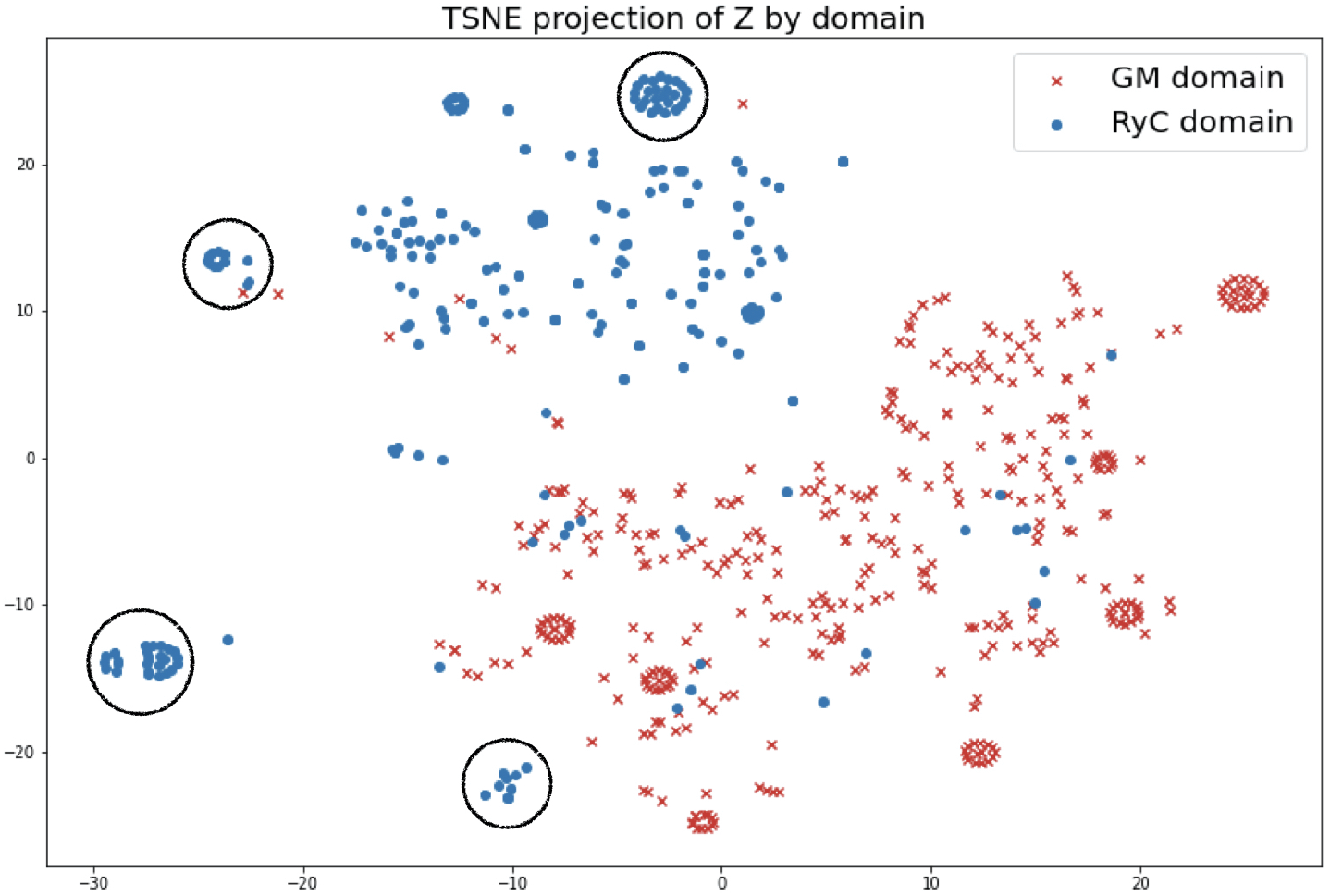
t-SNE 2-dimensional representation of the 14 latent variables of **Z** that are relevant for the domain view. Red crosses stand for a **z**_n,:_ whose observation comes from the GM domain, while every blue dot stands for a **z**_n,:_ whose observation comes from the RyC domain.

KSSHIBA can reduce the dimensionality from an initial MALDI-TOF of 10,000 features to only 76 latent features. This low-dimensional latent space representation together with the domain characterization justifies the better results of KSSHIBA over all SOTA approaches in the inter-domain scenario, as it can detect the differences between the MALDI-TOF depending on their domain learning to generalise better the prediction.

## 4. Conclusions

In this work, we predicted the AR of *K. pneumoniae* to ESBL and CP production based on both the MALDI-TOF spectra and their epidemiological information. In particular, we adapted the KSSHIBA algorithm to AR prediction outperforming, in terms of AUC, current SOTA algorithms such as XGBoost, LightGBM, MLP, SVMs, and GPs. At the same time, KSSHIBA’s application to MALDI-TOF is the first method that works with raw MALDI-TOF data, getting rid of external preprocessing with MQ. Moreover, it is the first method that provides dimensionality reduction by combining, simultaneously, MALDI-TOF and epidemiological data into a low-dimensional latent space. KSSHIBA also automatically adjust the model hyperparameters by Bayesian inference. Lastly, KSSHIBA provides explainable results by exploiting the epidemiological information, by means of its multiview architecture.

All these advantages are analysed over two different bacterial domains: (1) using data from the same hospital (GM), and (2) grouping strains from 18 geographically dispersed hospitals, selected by their phenotypic and genotypic resistance to beta-lactams (RyC). In Table 6 all current non-heterogenous models, such as GPs or SVMs, showed an overfit to one of the domains performing poorly in the smaller domain. Thus, heterogeneous models, which are able to analyse epidemiological information, are needed to predict AR fairly across domains. The experiments demonstrated that it is critical to adjust different distributions when working with two domains simultaneously. Thus, a linear kernel is needed to not overfit one domain. Due to that, the results conclude that a heterogeneous model with linear kernels, such as KSSHIBA LINEAR, has to be used to predict the AR susceptibility in *Klebsiella pneumoniae*. In fact, we have checked that the domain information improved the learning process by making KSSHIBA able to properly model the different data distributions by getting rid of the bias introduced by the data itself, which can lead to overfitting to one of the distributions, mainly if it presents domain unbalance. However, learning a probabilistic model entails learning a full distribution of the data which adds complexity to the training phase. Moreover, a double ARD prior to the kernel representation should be included in future work to be able to add feature selection to the model. Doing so, the model could be able to automatically select which peaks of the MALDI-TOF data are relevant for each prediction, which now, is a limitation.

Our contribution is, therefore, a step forward towards the goal of reducing ineffective antibiotic prescribing by being able to predict possible resistance mechanisms in *K. pneumoniae*. Its implementation in microbiological laboratories could improve the detection of multidrug-resistant isolates, optimising the therapeutic decision and reducing the time to obtain resistance mechanism results in comparison to nowadays manual methods.

As future work, we propose a longitudinal study over real test samples in clinics where the KSSHIBA model and other baselines are used to automatically predict the AR to check its viability in a real scenario. After that validation process, a web server would be implemented as a rapid AST method in laboratories.

## Acknowledgment

This work was supported by Spanish MINECO (Agencia Estatal de Investigación) [RTI2018-099655-B-10, PID2021-123182OB-I00, and TED2021-13182313-10 to P.O.]; this paper is also funded by MCIN/AEI/10.13039/ 5011000110 [PID2020-115363RB-I00 to A.G., C.S., and V.G]; and Comunidad de Madrid [IND2018/TIC-9649, Y2018/TCS-4705 to P. O.]; and the European Union (European Regional Development Fund and the European Research Council) through the European Union’s Horizon 2020 Research and Innovation Program [714161 to P. O.]; and Intramural Program of the Gregorio Marañón Health Research Institute to A. G.; and Health Research Fund (Instituto de Salud Carlos III. Plan Nacional de I+D+I 2013-2016) of the Carlos III Health Institute (ISCIII, Madrid, Spain) [PI15/01073, PI18/00997 to A. C. and B. R.] partially financed by the European Regional Development Fund (FEDER) ‘A way of making Europe’; and Health Research Fund Miguel Servet contract [CPII19/00002 to B. R.]

We acknowledge the SUPERIOR Study Group which provided the RyC domain data and includes the following members: Antonio Oliver and Xavier Mulet (Hospital Universitario Son Espases, Palma, Spain); Emilia Cercenado (Hospital General Universitario Gregorio Marañón, Madrid, Spain); Germán Bou and M. Carmen Fernández (Hospital Universitario A Coruña, A Coruña, Spain); Álvaro Pascual and Mercedes Delgado-Valverde (Hospital Universitario Virgen Macarena, Sevilla, Spain); Concepción Gimeno and Nuria Tormo (Consorcio Hospital General Universitario de Valencia, Valencia, Spain); Jorge Calvo, Jesús Rodríguez-Lozano and Ana Ávila Alonso (Hospital Universitario Marqués de Valdecilla, Santander, Spain); Jordi Vila, Francesc Marco and Cristina Pitart (Hospital Clínic, Barcelona, Spain); and María García del Castillo, Sergio García-Fernández, Marta Hernández-García, Marta Tato and Rafael Cantón (Hospital Universitario Ramón y Cajal, Madrid, Spain). This study was sponsored by MSD Spain. The STEP Study Group includes the following members: José Melo-Cristino (Serviço de Microbiologia Centro Hospitalar Lisboa Norte, Lisboa, Portugal); Margarida F. Pinto, Cristina Marcelo, Helena Peres, Isabel Lourenço, Isabel Peres, João Marques, Odete Chantre and Teresa Pina (Laboratório de Microbiologia, Serviço de Patologia Clínica, Centro Hospitalar Universitário Lisboa Central, Lisboa, Portugal); Elsa Gonçalves and Cristina Toscano (Laboratório de Microbiologia Clínica Centro Hospitalar de Lisboa Ocidental, Lisboa, Portugal); Valquíria Alves (Serviço de Microbiologia, Unidade Local de Saúde de Matosinhos, Matosinhos, Portugal); Manuela Ribeiro, Eliana Costa and Ana Raquel Vieira (Serviço Patologia Clínica, Centro Hospitalar Universitário São João, Porto, Portugal); Sónia Ferreira, Raquel Diaz and Elmano Ramalheira (Serviço Patologia Clínica, Hospital Infante Dom Pedro, Aveiro, Portugal); Sandra Schäfer, Luísa Tancredo and Luísa Sancho (Serviço de Patologia Clínica, Fernando Fonseca, Amadora, Portugal); Ana Rodrigues and José Diogo (Serviço de Microbiologia, Hospital Garcia de Orta, Almada, Portugal); Rui Ferreira (Serviço de Patologia Clínica—Microbiologia—CHUA—Unidade de Portimão, Portugal); Helena Ramos, Tânia Silva and Daniela Silva (Serviço de Microbiologia, Centro Hospitalar Universitário do Porto, Porto, Portugal); Catarina Chaves, Carolina Queiroz and Altair Nabiev (Serviço de Microbiologia, Centro Hospitalar Universitário de Coimbra, Coimbra, Portugal); Leonor Pássaro, Laura Paixao, João Romano and Carolina Moura (MSD Portugal, Paço de Arcos, Portugal).

## Footnotes

1 Icons images were provided by smart.servier.com

2 https://github.com/aguerrerolopez/RMPrediction

